# Short and long-term consequences of Habitat Transformation for Biodiversity

**DOI:** 10.64898/2026.04.21.719287

**Authors:** Brennen Fagan, Inês S. Martins, Jon W. Pitchford, Susan Stepney, Chris D. Thomas

## Abstract

Humans have dramatically altered the Earth’s surface and the distributions of species, but coherent patterns distinguishing cause from effect are hard to discern. Community composition and ecosystem function can change as rapidly as local land-use, and we lack robust principles to understand the outcomes of these changes. This hampers assessments of ecosystem collapse or robustness. Here we create theoretical ecosystems using a community assembly framework in which we manipulate environmental filtering via both the local land-use and species traits. This isolates the impacts of environmental filtering into land-use, land-use change, and species trait diversity, allowing us to extract clear patterns and relationships. First, we identify the paradox of maladaptation. We find that better land-use adaptation reduces species richness in the habitat but increases species abundance and ecosystem complexity. Increasing diversity amongst species traits reduces species richness via a similar mechanism. Additionally, whilst over long time scales there is very little effect of land-use change, on very short time scales there is a predominantly negative effect on richness. Together, this highlights the need for careful facilitation and management of land-use change in the face of an ever-changing world.

**Author Summary:** Humans have dramatically altered the Earth’s surface, primarily through land-use change and changing where species are and where they can go. This has resulted in extinctions, immigration, and, most of all, variation in where species end up. Here, we use simulations to compare how habitat, habitat change, and species’ adaptations create variation in outcomes due to land-use change. First, we show that while adaptation is good for individual species, it also reduces the overall diversity (number of species) of the ecosystem; similarly, well-adapted species out-perform and exclude less well-adapted species, also lowering the resulting diversity of the ecosystem. Secondly, we find that habitat change has different short- and long-term consequences. Habitat change can create dramatic declines on short time scales due to the different pressures experienced by species based on which species they eat (trophic level). But, on long time scales it has very little effect on diversity, as communities acquire new species adapted to the new conditions. We conclude that short-term declines in diversity due to recent or ongoing habitat change could be eased by helping well-adapted species to immigrate to form ecological communities in the new environment.

## Introduction

Humans are responsible for altering much, if not all, of the Earth’s surface [1]. These changes are widely credited with reducing biodiversity levels from local to global scales, fuelling concerns that the planet is entering a major extinction event ([2], but see [3]), and a degradation of the goods and services associated with relatively ‘intact’ ecosystems [4]. However, while global-scale research typically focuses on the average trends (and average differences between more and less modified environments), nearly all studies reveal a second phenomenon: variation. Variation among individual studies and localities, among regions and ecosystems, and across the different types of metrics used and data analysed can be so great that it is difficult to detect general trends [5]. Understanding the basis of this variation is a critical component of understanding human influences on the biosphere and, potentially, in developing future interventions to mitigate against potential losses.

The greatest local impacts on terrestrial diversity, and greatest projected threats, are thought to relate to anthropogenically-driven land-use changes [4]. Comparisons of human-modified environments with ‘relatively pristine’ ecosystems frequently reveal human-associated reductions in local species richness [e.g., 6,7]. Yet this trend is not universal, with several studies reporting opposite trends across various locations and ecosystems [e.g., 8], often linked to increased environmental heterogeneity through time. For instance, farming practices over the Holocene likely created a patchwork of anthromes intermixed with existing ecosystems, mostly via forest clearing [9]. These processes not only allowed for a diverse set of coloniser plants naturally associated with open and disturbed ground to take root in some places, but most likely generated scale-dependent ecosystem richness and diversity patterns [10]. The composition of past, present and possible future local ecological communities is a result, therefore, of the nature of the new environments that have been created (type of habitat), the availability of potential colonists (regional species pool), and their particular functional traits and adaptations (habitat associated performance differences) to the different habitat types. How these dimensions interact to shape current patterns of local biodiversity change remains poorly understood. Moreover, it is not clear how local communities may behave and re-structure before, during and after land-use change, because appropriate temporal comparisons in such locations (i.e., locations’ ongoing different levels of land-use change) require long-term high-resolution monitoring data, and thus are rare to find in the literature.

Here, we take a modelling approach to help understand the circumstances that can lead to either temporal decreases or increases in local species richness among functionally diverse ecological communities experiencing sudden habitat changes. Specifically, we apply the model (Figure 1a) developed by Fagan et al. [11] to explore how the diversity, dynamics and composition (‘basal’ versus ‘consumer’ species; cf. plants and animals) of the resulting ecological communities depend on (i) the habitat adaptations (performance differences associated with different habitat types) of species available to immigrate from the regional species pool (Figure 1b), (ii) the habitat type present in the patch (Figure 1c, Initial habitat type), and (iii) changes in the patch habitat types half way through simulation runs, as a representation of land-use change (hereafter referred to as *interventions*) (Figure 1c, Level of intervention).

**Figure 1:**
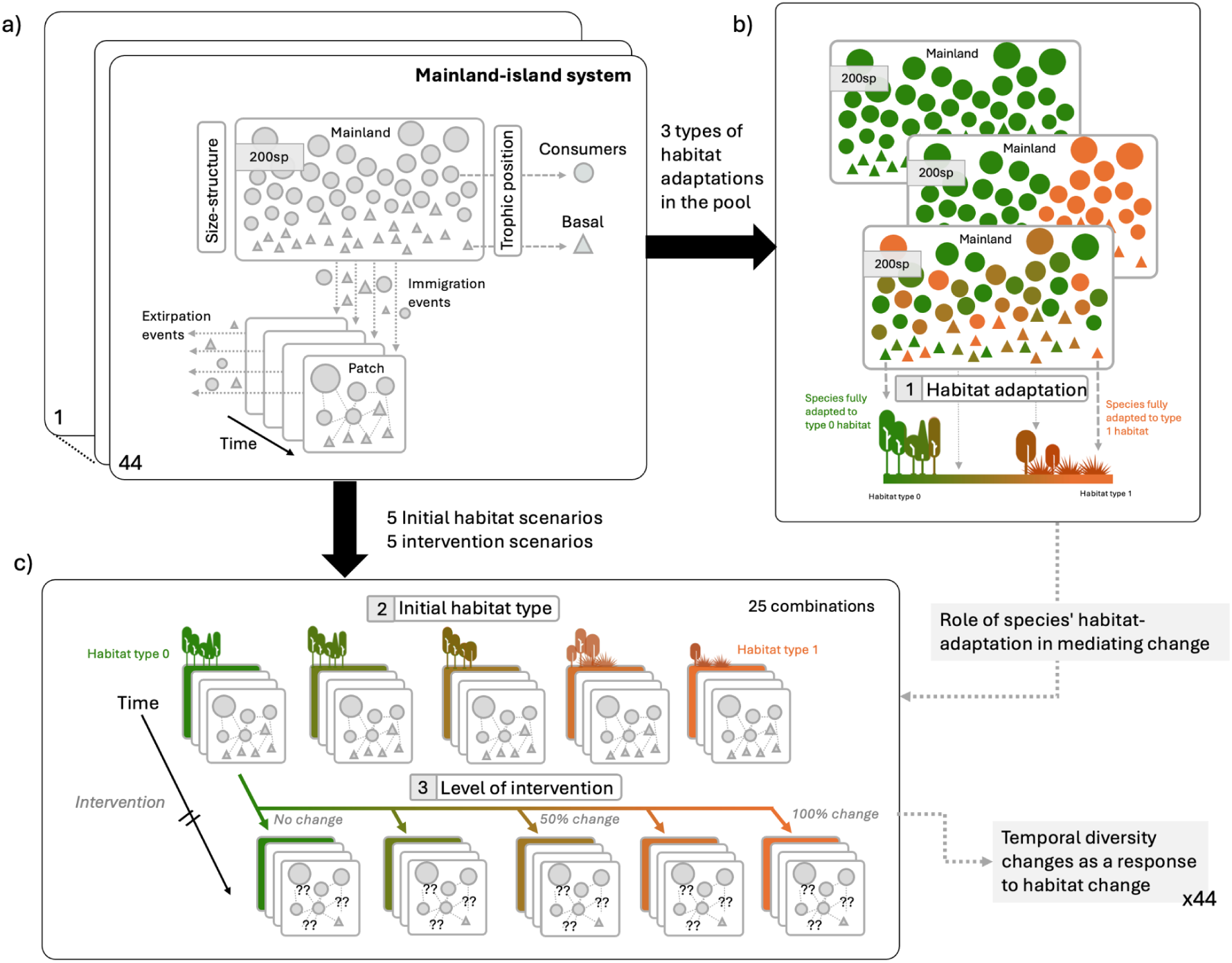
Conceptual diagram illustrating the different dimensions of our study design. **a)** Each simulated system (n = 44 replicates) includes a pool of 200 species and a single habitat patch. Each icon represents a species, while different shapes represent their trophic position (triangles = basal species, circles = consumer species; larger symbols indicate species with greater body size); the habitat in each simulation run is also continuously exposed to a series of stochastic immigration and extirpation events. Within each patch, species have predator-prey dynamics. Changes in diversity through time as a response to land-use change (intervention) can depend on **b)** the habitat adaptation of the species in the available pool of immigrants (different colours representing different habitat associated performance); and **c)** the specific type of intervention. Across our model runs we varied both the pool characteristics (3 distinct scenarios of possible habitat adaptations), the initial habitat type (5 scenarios) and the level of intervention (5 scenarios). We track community composition and diversity (taxonomic and functional) change within the patch.

The habitat types and adaptations both range from 0 to 1, such that species adapted to type 0 have peak performance (maximum net growth) in habitat type 0, etc. Habitat interventions in the patch can range from modest (e.g., type 0 transitions to type 0.25) to major (e.g., habitat type 0 transitions to type 1), with no change as intervention ‘controls’ (e.g., type 0 throughout) (see Methods for further details).

To demonstrate how different diversity trajectories can occur under various environmental conditions, we present results across key scenarios of change. First, we show the expected diversity and dynamics of communities under a constant environment – where all species in the regional pool share the same habitat adaptations and the habitat type stays the same (i.e., no intervention). We then investigate how interventions change the observed communities, separating out the effects over different time scales. Afterwards, we study how diversifying the adaptations present in the pool, first without and then with interventions, change the diversity of the resulting communities. The results show that the species richness and functional diversity of model communities can either increase or decrease (sometimes in non-intuitive ways) as a consequence of the matches and mismatches between species habitat adaptations in the pool and the nature and degree of intervention. In summary, this modelling strategy allows the roles of species adaptations and land use change to be disentangled across a range of ecological and land-use change scenarios.

## Results

### Diversity when all species share the same habitat adaptation

When all species in the regional pool have the same habitat adaptation (all species have maximum performance when in habitat type 0 in Figure 2), and no interventions are imposed, the simulation results show that ‘poorly adapted communities’ in individual habitat patches achieve higher richness than ‘well adapted communities’ (Figure 2a, b). Specifically, richness tends to be lowest when species are highly adapted to the habitat patch (species type 0 in habitat type 0; cyan line in Figure 2a), and highest when species are poorly adapted to it (species type 0 in habitat type 1; yellow line in Figure 2a). Note that low diversity is different to low abundance; Figure 2c shows that abundances are generally high in well adapted low diversity communities. These trade-offs are driven by changes at species level; basal species are better able to persist in habitats where they are poorly adapted compared to consumer species (Figure 2d, e right, Figure S1), generating the highest basal diversity in poorly-adapted communities (Figure 2d, examples shown in Figure 2a insets). Consumer richness remains relatively constant, although their abundance goes down, until all species are poorly adapted (species type 0 in habitat type 1), when consumer richness and abundance become very low (Figure 2d, e). As a consequence, maximum richness occurs when the habitat type is still relatively inhospitable (species type 0 in habitat type 0.75; red), but not so bad that consumer diversity collapses (species type 0 in habitat type 1; yellow) (Figures 2b, d). Because these simulations control for other sources of variation, this pattern in diversity emerges as the product of the species adaptation-habitat type (mis-)match influencing species-interactions and food web structures and dynamics (see below and Figure S2 for sensitivity to interaction strength). In summary, in the absence of habitat change, poorly adapted communities have the highest richness.

**Figure 2:**
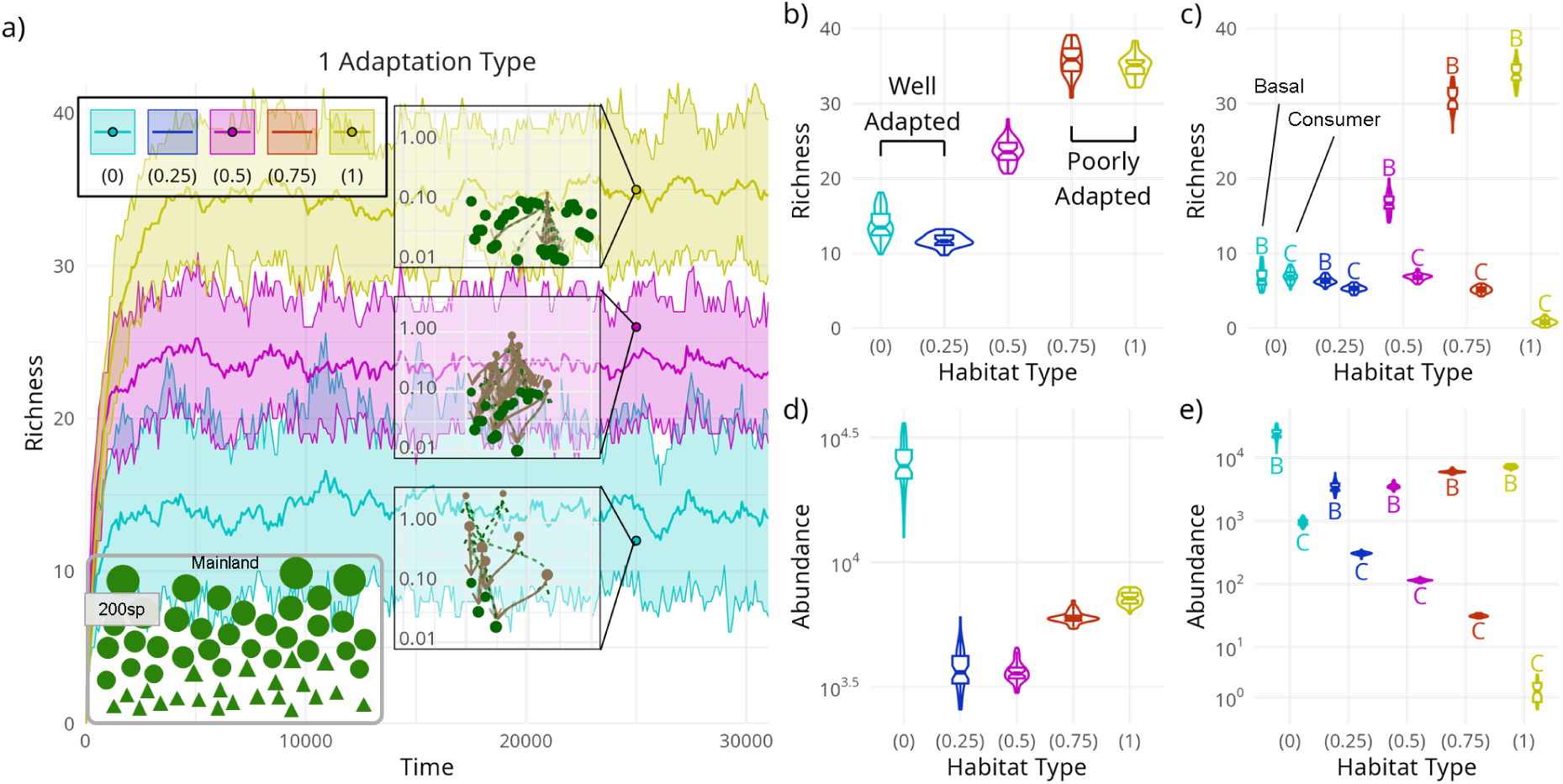
Emerging trade-offs between diversity and habitat type. a) Temporal changes in species richness across 44 systems for 3 types of habitat (intermediate scenarios are omitted for simplicity) in the absence of habitat interventions. Different colours represent different habitat scenarios (e.g. yellow: the habitat is of type 1). For a given scenario, solid lines show the mean across the 44 system runs, while ribbons show the range containing 75% of the richness values across the systems through time. Points represent one system evaluated at the same time for each scenario. Insets show how resultant communities, points in a, are structured for the different scenarios (brown = consumer species, green = basal species, with lines connecting consumer and consumed species). b) Violin plots of richness averaged across times 20,000 and 30,000 for each simulation as a function of habitat type. Note that species adaptation to the habitat is highest in the left hand side scenarios, and lowest in the right hand ones. c) As in panel b, but with richness displayed by whether the species are basal species (violins marked B) or consumer species (violins marked C). d-e) As in panels b and c, but for total abundance (i.e., abundance summed over all species in panel d or over all basal or consumer species in panel e).

We then examined the effect of interventions on biodiversity, i.e., we changed the habitat type of the patches half way through the simulations to evaluate the effect of habitat change. As above, adaptation for habitat type 0 was common across all species. Over long time scales, the level of intervention and starting habitat type exhibited minimal influence on final ecosystem composition; instead the community structure and ’final species richness’ after intervention (Figure 3a) closely matched those of no-intervention runs with the same final habitat type (Figure 2a).. For instance, interventions changing habitat type 0.5 to habitat type 1 showed a long-term increase in diversity, with richness levels similar to the ones documented for a no-intervention habitat of type 1 (compare the final state of the red line in Figure 3a with the yellow line in Figure 2a). Overall, and despite some level of variation among the 44 systems and intensity of perturbation, no overall measurable long term effect could be detected.

**Figure 3:**
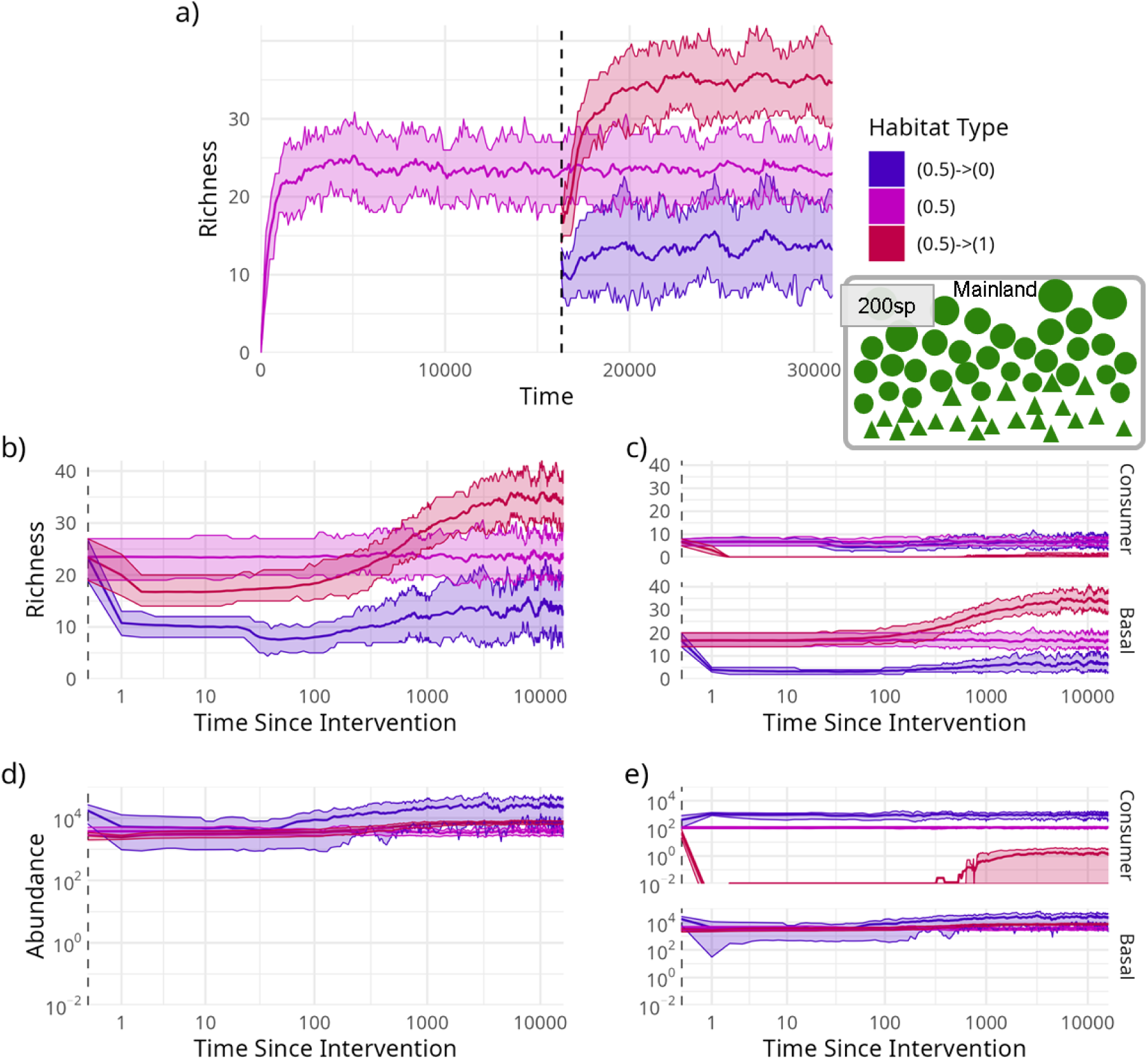
Interventions on long time scales produce little to no effect, but can create dramatic changes on short time scales. **a)** Temporal changes in species richness across 44 systems for 3 habitat change scenarios: intermediate 0.5 habitat type without intervention and with intervention to either extreme 0 or 1 habitat types. The vertical dashed black line indicates when interventions occur. **b)** Richness changes after the (non-)intervention on a log(1+Time) axis highlighting the difference in short term and long term post-intervention behaviour. **c)** As in panel b, but with richness partitioned amongst consumer (top panel) and basal (bottom panel) species. **d)** As in b, but for total abundance. **e)** As in c, but for total abundance. For all panels, solid lines show the mean across the 44 system runs, while ribbons show the range containing 75% of the richness values across the systems through time.

In contrast to the lack of a long-term effect, the short-term effect of intervention was to generate a decline in species richness (Figure 3b) during the initial transition period, regardless of whether those communities would decline or increase in the longer-term. Transitions from a habitat type that species were well-adapted to to a poorer match caused losses in consumer species (Figure 3c), driven by simultaneous reduced resource availability (initial decreases in prey abundance before recovery via the colonisation of additional basal species from the pool) and increased mortality (direct effect of adaptation, Figure 3d, e). In contrast, going from less preferred to a more preferred habitat type generated losses of basal species (Figure 3c), as consumer species’ total abundance increased (Figure 3e) and consumed many basal species to extinction. These effects generally last for ∼1000 time units before approaching steady-state values (Figure 3b, d).

Figure 4 summarises these findings across the full range of interventions. Each cell shows the % change in richness (top) and abundance (bottom) caused by interventions at short (left) and long (right) time scales, so that the top-left and bottom-right cells show situations where interventions are large, in contrast to the bottom-left to top-right diagonal which represent no change. While short term effects are due to a combination of initial and final habitat types (Figure 4 left), long term effects only depend on the final habitat type (Figure 4 right, SI Figure S3). For instance, communities well-adapted to type 0 habitats in patches that transitioned from sub-optimal habitat types to habitat type 0 (bottom-right quadrant in the top-right panel of Figure 4) exhibit the highest long-term declines in richness, with losses up to 65.5%, mostly due to basal species extirpations as consumer species thrive (SI Figure S4). Overall, communities eventually acquired the ‘expected diversity’ and consumer-basal composition of the ‘destination’ environment, but consistently exhibited a diversity ‘trough’ during the transition phase (SI Figure S3).

**Figure 4:**
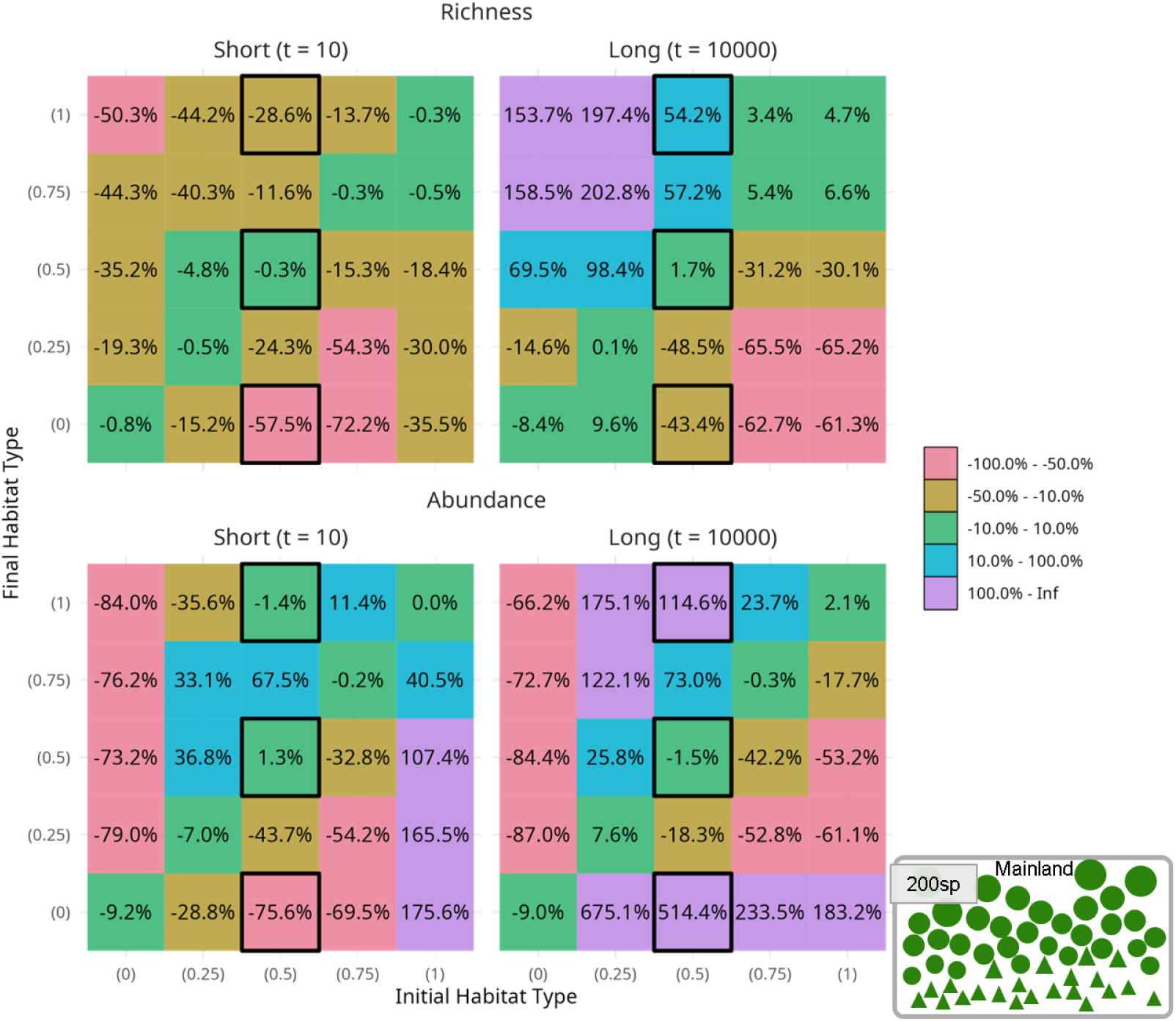
Contrasts between initial and final habitat types vary through time when there is a single adaptation type. Heatmaps show average percentage change (see Methods) across the 44 systems for each combination of initial and final habitat types for richness (above) and abundance (below) across short (left, 10 time units after intervention) and long (right, 10000 time units after intervention) time scales. The short time scale analysis compares the community at 10 time units after intervention to the community the moment just before intervention, while the long time scale analysis compares the average across 9500 - 10500 time units after intervention to the average 1000 time units before intervention. Black boxes indicate scenarios shown in Figure 3. For a breakdown between basal and consumer species see SI Figure S4.

### Diversity when species vary in their habitat adaptations

Given the importance of the relationship between species adaptations and habitat type, we also considered the dynamics and structure of communities where the adaptations in the species pool were either mixed (50% of species adapted to habitat 0, 50% adapted to habitat 1, hereafter “2 Adaptation Types”), or contained a range of adaptations (with habitat adaptation values drawn from a uniform distribution between 0 and 1, hereafter “Multiple Adaptation Types”). This is in contrast to every species being adapted to habitat type 0 in Figures 2-4.

In the absence of habitat interventions (Figure 5), simulations show that the long-term effects of the environment on richness are negligible (richness is approximately the same for different habitat types), with one notable exception. That was when the pool was constructed of species with 2 adaptation types while the patch had habitat type 0.5 (Figure 5a). This was the one situation in which all species were equally maladapted, resulting in greatly increased species richness (again driven by basal increase; SI Figure S5). For all other scenarios, the better adapted species (i.e., those with a closer match between their habitat adaptation and the habitat type of the patch) outperformed others (Insets in Figure 5a, d) and hence formed a majority of species in the community (with richness similar to species with a single adaptation type in type 0 habitat, Figure 2). For instance, when species have multiple adaptation types, species with adaptations of approximately 0.25 predominate in 0.25 habitats, 0.5 species in 0.5 habitats, and so on - the asymmetry in Figure 5d (Insets) is generated by the fact that species adaptations in the pool are bounded by 0 and 1. The outcomes are again modulated by the balance between basal and consumer species, following the patterns established in Figure 2.

**Figure 5:**
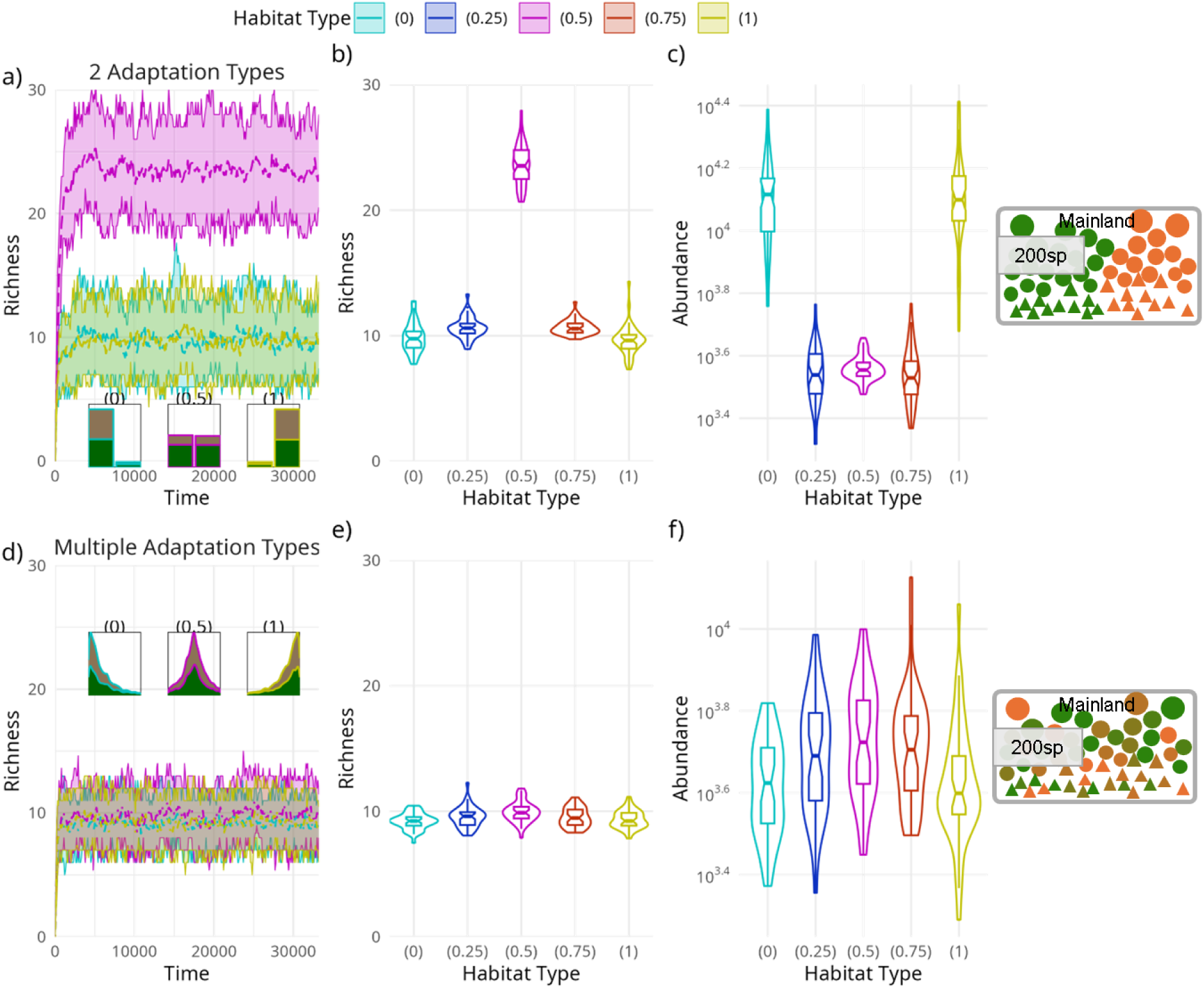
Diversity of habitat type adaptations reduces emergent richness in the local ecosystem. The first row shows results for scenarios with 2 adaptation types in the pool: half of species with adaptations to habitat type 0, half with adaptations to habitat type 1; the second row shows results for scenarios with multiple adaptation types in the pool: species adaptations range from 0 to 1. **a)** Temporal changes in species richness across 44 systems for 3 types of habitat, with lines showing the mean and ribbons the 75% interval of richness values through time (compare Figure 2a). Inset histograms capture the distribution of (either-or) habitat adaptation traits and species type (basal, green or consumer, brown) of the emergent communities. **b)** Violin plots of richness averaged across times 20,000 and 30,000 for each simulation as a function of habitat type (compare Figure 2b). **c)** As in panel b, but for total abundance on a logarithmic scale (compare Figure 2d). **d-f)** As in a-c), but for Multiple Adaptation Types. Insets are stacked kernel density estimates instead.

The long-term effects of habitat intervention in scenarios with multiple adaptation types were as expected - species richness did not substantially change because the richness associated with the initial habitat type and that of the final habitat type were comparable (Figure 6a; for scenarios with 2 adaptations types see SI Figure S6). Notably, transitions still consistently generated a short-term decline in richness as species adapted to the previous habitat type were replaced by those adapted to the new habitat type, especially when transitions were from intermediate to extreme habitats (Figure 6b, e). Despite the relatively small changes in richness, these transitions often caused pronounced shifts in community structure (SI Figure S7).

**Figure 6:**
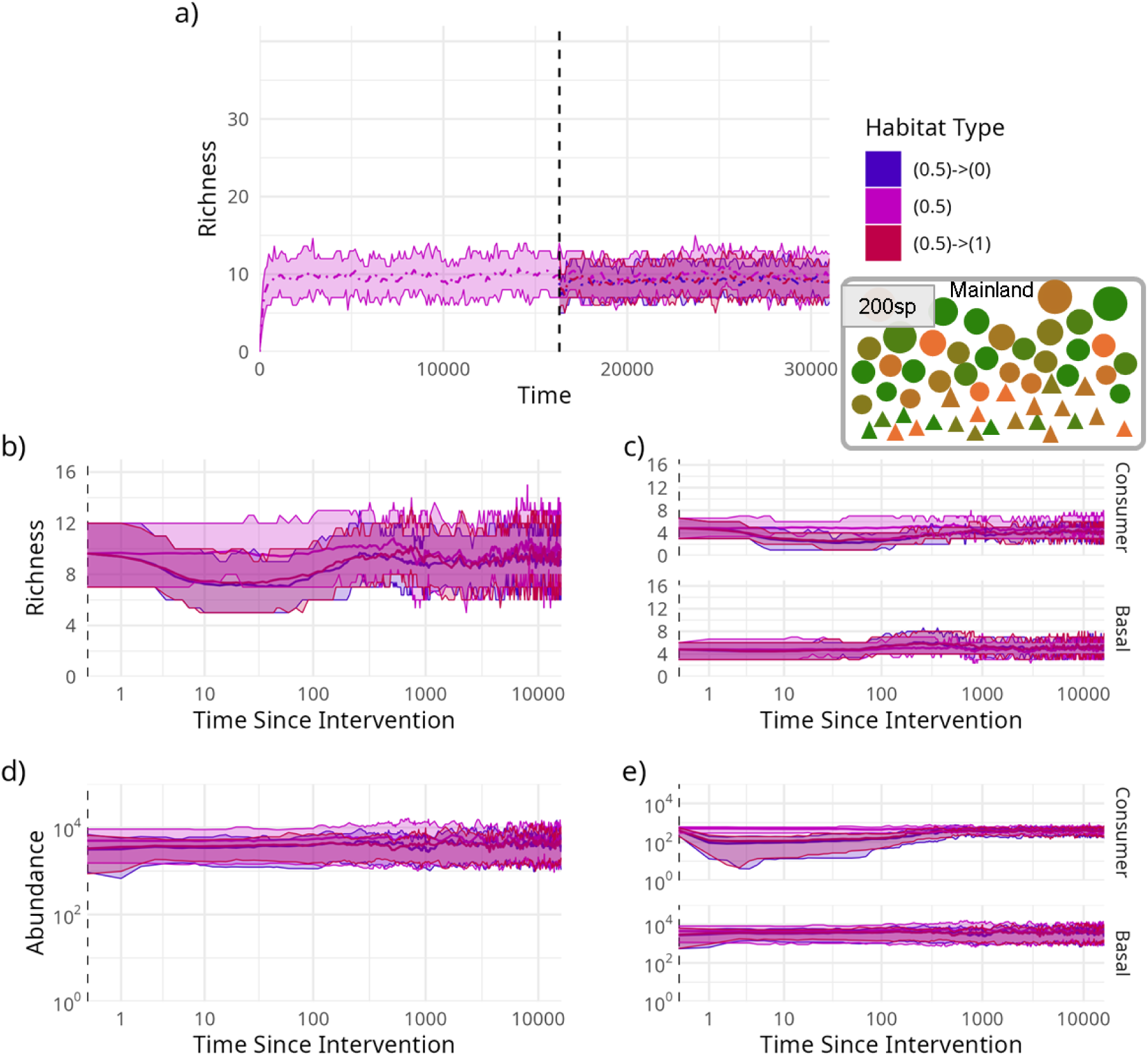
Diversity of habitat type adaptations in the pool obscure changes in the structure of the system over short time scales. **a)** Temporal changes in species richness across 44 systems for 3 habitat change scenarios: intermediate 0.5 habitat type without intervention and with intervention to either extreme 0 or 1 habitat types. The vertical dashed black line indicates when interventions occur. **b)** Richness changes after the (non-)intervention on a log(1+Time) axis highlighting the difference in short term and long term post-intervention behaviour. **c)** As in panel b, but with richness partitioned amongst consumer (top panel) and basal (bottom panel) species. **d)** As in b, but for total abundance. **e)** As in c, but for total abundance. For all panels, solid lines show the mean across the 44 system runs, while ribbons show the range containing 75% of the richness values across the systems through time.

Restructuring occurs first at the higher trophic levels: large-bodied consumer species are lost, similar to transitions to habitat types to which communities are poorly adapted (Figure 3c), before being replaced by smaller consumer species favoring the new habitat type (Figure 6c, SI Figure S7). This is followed by gradual changes at lower trophic levels: basal species adapted to the new habitat become more abundant and frequent (Figure 6d), followed by the recovery of larger consumer species better adapted to the new habitat type. This combines elements of transitions observed when the regional pool had only one type of habitat adaptation (Figure 3).

Across the full set of intervention scenarios, the short term response to habitat change was characterized by a predominant decline in species richness (Figure 7, top-left). Short-term declines in richness, particularly in the most extreme habitat interventions, reflect the onset of local species extirpations following the rapid shifts in habitat suitability, and the slower pace of colonization by new (and better adapted) immigrants from the pool (SI Figure S7). However, these biodiversity declines are transient across the simulations, and as time passes (t = 10000), the richness changes become close to zero (Figure 7, top-right). The changes in richness are somewhat decoupled from change in abundance - while richness seems to return to pre-intervention levels, total abundance often shows long-term changes (Figure 7, bottom-right), particularly in transitions involving intermediate habitat types.

**Figure 7:**
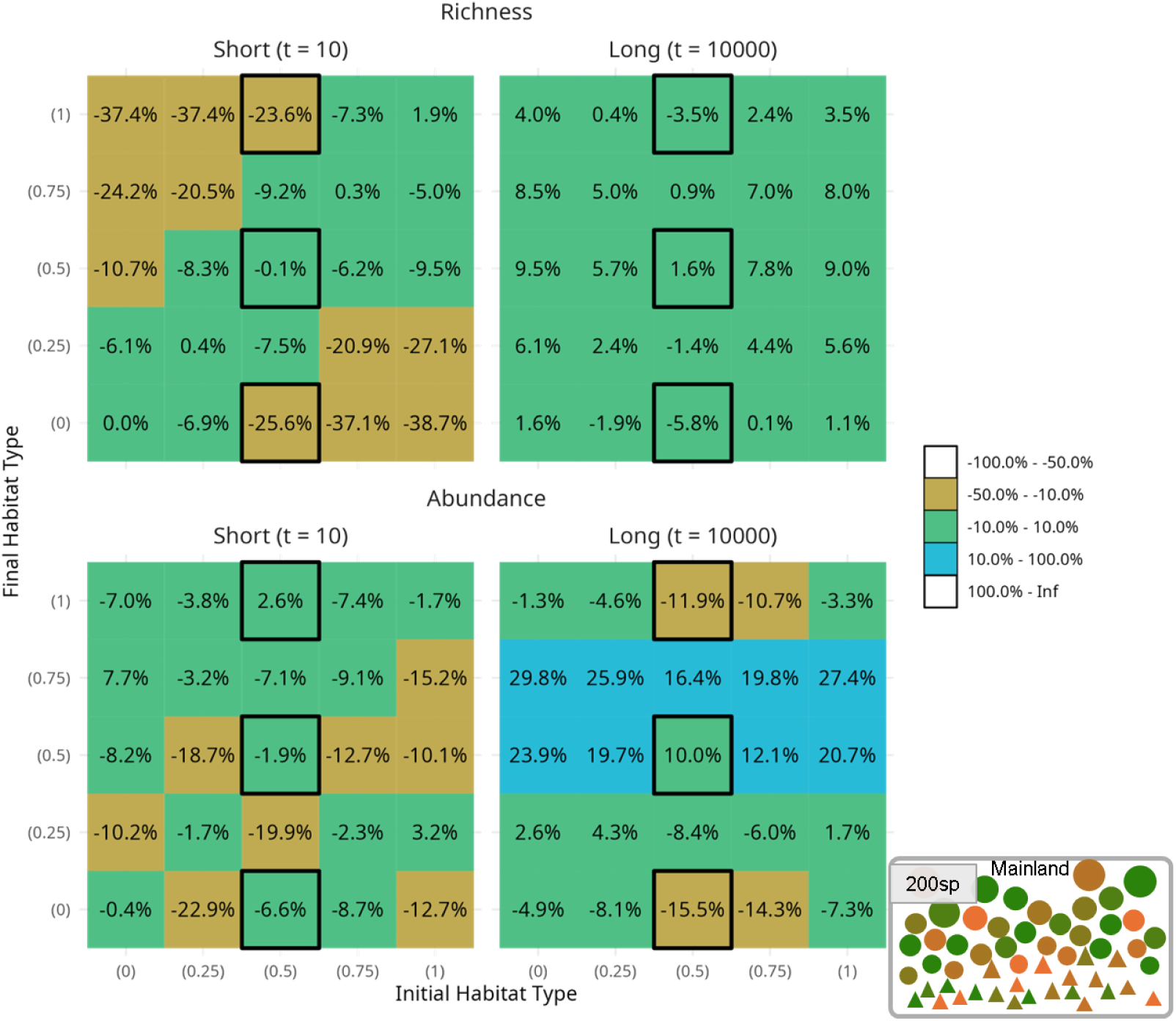
Contrasts between initial and final habitat types vary through time when there are multiple adaptation types (compare Figure 4). Heatmaps show average percentage change (see Methods) across the 44 systems for each combination of initial and final habitat types for richness (above) and abundance (below) across short (left, 10 time units after intervention) and long (right, 10000 time units after intervention) time scales. The short time scale analysis compares the community at 10 time units after intervention to the community the moment just before intervention, while the long time scale analysis compares the average across 9500 - 10500 time units after intervention to the average 1000 time units before intervention. Black boxes indicate scenarios shown in Figure 6. For a breakdown between basal and consumer species see SI Figure S8.

While losses are temporary, we also observe that, in comparison to reference richness baselines just before the intervention takes effect, our systems have stochastic fluctuations (due to species extirpations and immigrations) even across the scenarios without intervention (SI Figure S9). Larger interventions result in more simulations with rapidly decreasing richness (0 to/from 1, SI Figure S9), while smaller changes tend to cause systems to decrease slightly later (0 or 1 to/from 0.5, SI Figure S9; note the slightly higher richness of 0.5 reduces the impact, see Figure 5b, bottom).

## Discussion

### Connecting complexity, stability and maladaptation-richness paradox

We have shown that adaptation to the environment, in the form of increased reproduction or decreased mortality, generally decreases ecosystem richness. Changes in abundance due to adaptation drive these changes in richness in two different ways: poorly adapted species are more easily displaced by better adapted species, and, since the interaction terms scale with species abundance, better adapted species also have stronger de facto interactions. Stronger interactions contribute to instability, consistent with established theory connecting complexity and stability [12,13] and with the proposed maladaptation-richness paradox [14]. The richest ecosystems in our simulations, however, are neither functionally diverse nor ecologically interesting – consisting almost entirely of basal species with almost entirely the same traits – lending to a zoo-like structure with species neither abundant nor interacting. The alternative, an abundant and complex but species-poor and unstable community, results from a pool of species of whom at least some are very well-adapted, possibly reflecting an increasingly connected world.

### Understanding short term biodiversity loss

Over very long time scales the consequences of land-use change may be negligible. Over short time scales, immediately after changes in land-use, the ecosystem rapidly reorganises. Local abundances respond faster than immigration-driven processes, leading to a variety of consequences. First, the majority of – but not all – transitions following land-use change result in declines in richness, although the nature of the declines depends on the diversity of adaptation to the new environment. Declines are particularly prominent when land-use changes are larger, and when the species have very limited land-use preferences. When the land-use changes are small and the regional pool contains species with a larger variety of preferences, the change in richness is more likely to be slight or positive and to be perceived as a part of the natural succession of the ecosystem.

### Caveats and future directions

While our model allows for assembly of a community that adapts to changing land-use, our model does not incorporate evolution. This allows us to pinpoint the effects of land-use and land-use change in our simulations, but it neglects a key component of ecosystem resilience in doing so. Incorporating evolution, both in the ecosystem as well as in the regional pool, has contrasting arguments for whether it would increase or decrease the resilience of the ecosystem [15,16]. The software framework presented here, which is publicly available, could be adapted to explore these issues. However, doing so would involve making modelling decisions about eco-evolutionary time scales which are outwith the scope of this study.

Similarly, we focus on a single, essentially disconnected habitat, an assumption which can also be relaxed by adapting our software framework [11]. Incorporating more patches would allow for complications such as reservoir populations and source-sink dynamics beyond just the inclusion of heterogeneous landscapes. These effects would likely obscure the effects of land-use and land-use change in our simulations, including their role in community assembly and environmental filtering, but are likely to be crucial in an increasingly connected world. Similarly, we assume that there were no correlations between land-use preference and the immigration (or extirpation) rates of species from the regional pool. This might be an important additional factor, but its inclusion would prevent us from exploring the full space of ecosystem responses to land-use, land-use change, and land-use preferences of species.

## Conclusion

We have used mathematical models and computer simulations to examine the effects of land-use adaptation and change over both long and short timescales. Our results reflect the need to accommodate an increasingly well-connected world in which species can, and will, find their preferred land-use, emphasising that this will likely lead to higher turnover and more immediate losses than gains. These new ecosystems are more likely to have a smaller number of more abundant species, but these new ecosystems are also likely to have more complex interactions and more dynamic change. Interpreting these trends in the context of effective management and biodiversity conservation remains a global challenge.

## Materials and Methods

### Model

Our methodology is a direct expansion of a size-structured, consumer-basal generalised Lotka-Volterra mainland-island model with temporal coupling between community dynamics, immigration and extirpation events [11]. This model was itself an iteration on a community assembly model used previously to study community assembly over long periods of time [17] and to assess the role of assembly graphs in population cycles [18]. More similar to the original version of the model, we restrict ourselves to a single patch without patch-to-patch dispersal to focus on the effects of land-use change (interventions).

We modified that original model to include an additional trait for each species denoting its habitat adaptation, used for determining a species’s performance in a given habitat. Land-use in the patch (the patch’s “trait”, or habitat type) and species adaptation (the species trait) are numbers in [0, 1] and are used to determine the effect on a species’s intrinsic reproduction or mortality due to being in a patch with a specific habitat type. These values are set at the beginning of a simulation and the species adaptations are not varied within a simulation. When habitat type is varied during a simulation (i.e., an intervention occurs), it is varied deterministically. In order to avoid direct, 3-way interaction effects between multiple species and the habitat, we chose to have the patch-species interaction affect the species’ intrinsic growth rate (if a basal species) or mortality rate (if a consumer species) as follows. We write

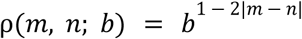

and

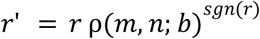

where ρ is the function which measures the effect of the difference between habitat type (𝑚) and species’ adaptation (𝑛) for a given intensity (𝑏, set to 5 in the main text), 𝑟 is the intrinsic growth or mortality rate of the species, 𝑠𝑔𝑛 is 1 if 𝑟 is positive (growth) and -1 if 𝑟 is negative (mortality), and 𝑟’ is the modified growth or mortality rate. We choose these functions so that a species in an extreme habitat type it is well-adapted to receives a five-fold multiplicative benefit, but if it is in the opposite habitat type, it receives up to a five-fold multiplicative penalty, regardless of whether it is a basal or consumer species. These changes directly modify the intrinsic 𝑟 species value in the previous versions of the model.

Additionally, we investigated the effect of the temporal coupling between the community dynamics and the immigration and extirpation events. This coupling was created by setting the rate of immigration opportunity events (as well as neutral extirpation events) to one-tenth the magnitude of the dominant eigenvalue, |λ_1_ |, of the interaction matrix, as the exact eigenvalues of the Jacobian that the community will experience cannot be known ahead of time and can vary substantially over the course of a simulation. This results in a per-system coupling between the events and the dynamics, allowing the events to occur more frequently (in absolute time) in systems with stronger interactions, so as to create consistent time scales across systems. To understand the robustness of this choice, we also compared the distribution of |λ_1_ | across the interaction matrices to those of the Jacobians during periods of little change post hoc; the standard deviation of |λ_1_ | across the interaction matrices was 0.0015, while for Jacobians it was 1.6, suggesting that the choice had comparatively little effect. We verified this by re-running the non-intervention simulations and re-analysing the results. We found no noticeable difference between results.

Similarly, we considered the effect of interspecific competition amongst the basal species. By default, our model includes predator-prey interactions between consumer species and their prey species which can induce indirect competition when consumer species are present, but some scenarios produce very few to no consumer species (Figure 2). We add this effect by setting a carrying capacity on the basal biomass, adjusting the intrinsic growth rate, re-running the non-intervention simulations, and re-analysing results. We found very little effect: in the 1 Adaptation Type scenario, this slightly altered the details of the pattern observed in the richness, but overall conclusions remain unchanged (SI Figures S10, S11).

### Simulations

To isolate the changes in our outputs to causal changes in species adaptations, habitat types, or changes in habitat types, we simulated 44 distinct pool-patch (mainland-island) metapopulations type systems (Figure 1). This number was chosen to give us a critical mass of replicates to make inferences upon while remaining computationally tractable. Each simulated system consists of a pool of 200 species (the source or mainland comprised of basal and consumer species with size-structured interactions [17]; Figure 1a), a single patch (the habitat or island), and a fixed, randomly generated “history” of events of immigrations and extirpations to be attempted (with varying success) over time [11]. Further extirpations can and will happen based on species interacting with each other and with the habitat itself (see below). Where time is used, we have rescaled the time by the characteristic eigenvalue of the interaction matrix, resulting in similar time scales for all simulations due to the coupling of the characteristic eigenvalue of the interaction matrix to the stochastic immigration and extirpation rates. Full simulation code is also available (see Data Availability).

For each system, we then tested how interactions between species and the habitat affects diversity and ecosystem structure by varying (1) the habitat-adaptation traits of species in the regional pool (mainland, source), (2) the initial habitat type of the patch, and (3) the level of intervention a patch experiences (land-use scenarios) (Figure 1). Specifically, and to test the effects of adaptation traits in mediating diversity across patches of different habitats, we created 3 distinct scenarios for regional pools (Figure 1b). First, we used a scenario where all species were assigned the same habitat adaptation trait value, so all species perform optimally in the same habitat type (in this case to habitat type 0), the 1 Adaptation Type scenario (Figure 2).

Second, we used a scenario where all species were evenly assigned one of two extreme values (0 or 1), so half the species perform best in habitat type 0 and the other half in habitat type 1, the 2 Adaptation Types scenario (Figure 5, top). Third, the species could be assigned values uniformly at random from the interval [0, 1], thus creating a pool characterized by species with varying optimal habitat types, the Multiple Adaptation Types scenario (Figure 5, bottom). For each pool type, we then explored different scenarios of habitat interventions (Figure 1c), which allowed us to test how diversity in the patch may change before, during and after different levels of habitat change. Here we explore 25 different scenarios, by allowing for 5 possible initial habitat types for the patch (0, 0.25, 0.5, 0.75, or 1), and 5 possible final habitat types (idem).

Note that when initial and final habitat type match, this corresponds to a no change scenario (5 out of the 25 scenarios). Where there is a difference in initial and final habitat types, we pick a time in the middle of the simulation (intervention point- calculated by averaging the median evaluation time and halfway to the last event, approximately 16,500 time units across simulations); at this time, the habitat type spontaneously shifts between the initial and final habitat types. Each simulated system is then run multiple times, varying just the habitat-type intervention scenarios. Because a given system is run multiple times with varying pool type and habitat-type intervention scenarios (i.e., 75 runs per system), our runs across systems are naturally paired so that their differences must be driven by the change in pool scenario or/and habitat-intervention scenario between any given two runs. Altogether, we carried out 3300 model runs. We then summarise the differences across sets of runs (organized by different types of scenarios) to create our results in the main text.

### Statistics

For descriptive statistics, we used common implementations provided by the ‘vegan’ package [19] in R [20], with data processing employing ‘dplyr’ [21], ‘foreach’ [22], ‘igraph’ [23], ‘tidygraph’ [24], and ‘tidytablè [25] packages. Plotting was accomplished with ‘cowplot’ [26], ‘ggraph’ [27], ‘ggplot2’ [28], and ‘ggpp’ [29].

For each time-point within a simulation run, we record the abundance of each species. When combined with the species traits for the given scenario, this allows us to construct richness (number of species) and total abundance (sum of abundance of species present) overall or for sub-sets of the species as well as the proportion of species that are basal or consumer or their adaptation types (as in Figure 5). We also record when species successfully colonise or when they extirpate during simulation runs, and we employ the interaction matrix and set of model equations to reconstruct food web networks.

Where we plot line plots, we plot the mean of the runs across the 44 systems as well as intervals containing 75% of the relevant statistic at the given point in time. We choose 75% for visualisation purposes only, i.e., to keep a sense of the variation while also avoiding overplotting. Where we evaluate statistics, unless otherwise specified, we evaluate the statistics between approximately 20,000 time units and 30,000 time units, which occurs beyond the time of any intervention. Keeping these numbers consistent between runs avoids complications regarding differences in sample or time series length. These times themselves are chosen to allow time for the post-intervention system to reach a new steady state. Evaluations of the underlying dynamical system, and thus the summary statistics, are not necessarily made at exactly these time points (but are made consistently at the same times across model runs of any given system in connection to the history assigned to each simulated system).

For the summary statistics presented in Figures 4 and 7, we proceed simulation run by simulation run. For short time scale comparisons, we initially compare the statistic at the moment just prior to the intervention (change in habitat type) to that 10 time units after the intervention. For long time scale comparisons, we instead compare the median statistic from the 1000 time units just prior to the intervention to the median statistic across 9500 - 10500 time units after intervention. In each case, we have 44 simulation run comparisons within a bin in Figures 4 and 7. To perform the comparison on a logarithmic scale – so that a multiplicative (e.g., 5-fold) increase is weighted evenly to a decrease – we calculate the comparison as log10(Post-Intervention Value / Pre-Intervention Value). To place this back on a regular, zero-centred scale, we average the logarithmic scale values before exponentiating, and then subtract 1. This results in a final formula of ((10^mean(log10(new value / old value)) - 1) *100%).

## Acknowledgements

We thank Jack Hatfield and the Macroscope Lab for helpful discussions. The Viking cluster was used during this project, which is a high performance compute facility provided by the University of York. We are grateful for computational support from the University of York, IT Services and the Research IT team. This work was funded by a Leverhulme Trust Research Centre – the Leverhulme Centre for Anthropocene Biodiversity – as a part of Leverhulme Trust Grant RC-2018-021.

## Data Availability Statement

All code, random seeds, R scripts, and summarised data used to generate the simulations and subsequent figures are available at https://github.com/Brennen-Fagan/Community-Assembly. The archived code and data can be found at ZENODO LINK.

## Supplemental Figures

**Figure S1:**
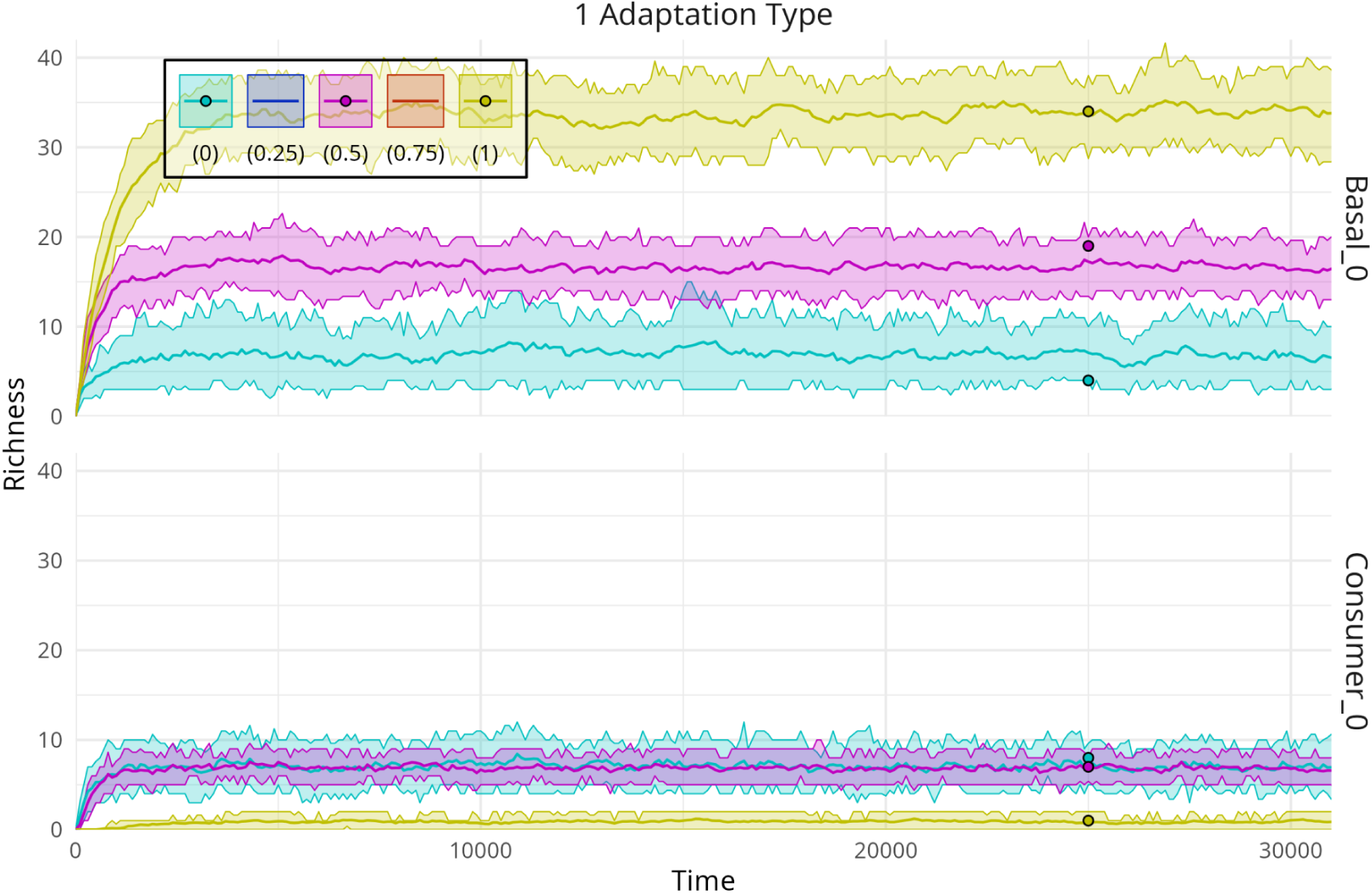
Emerging trade-offs between diversity and habitat type, broken up by basal-consumer species types. Temporal changes in species richness across 44 systems for 3 types of habitat (intermediate scenarios are omitted for simplicity) in the absence of habitat interventions. Different colours represent different habitat scenarios (e.g. yellow: the habitat is of type 1). For a given scenario, solid lines show the mean across the 44 system runs, while ribbons show the range containing 75% of the richness values across the systems through time. Points represent one system evaluated at the same time for each scenario. Above) Species richness due to basal species with adaptation to habitat type 0. Below) Species richness due to consumer species with adaptation to habitat type 0.

**Figure S2:**
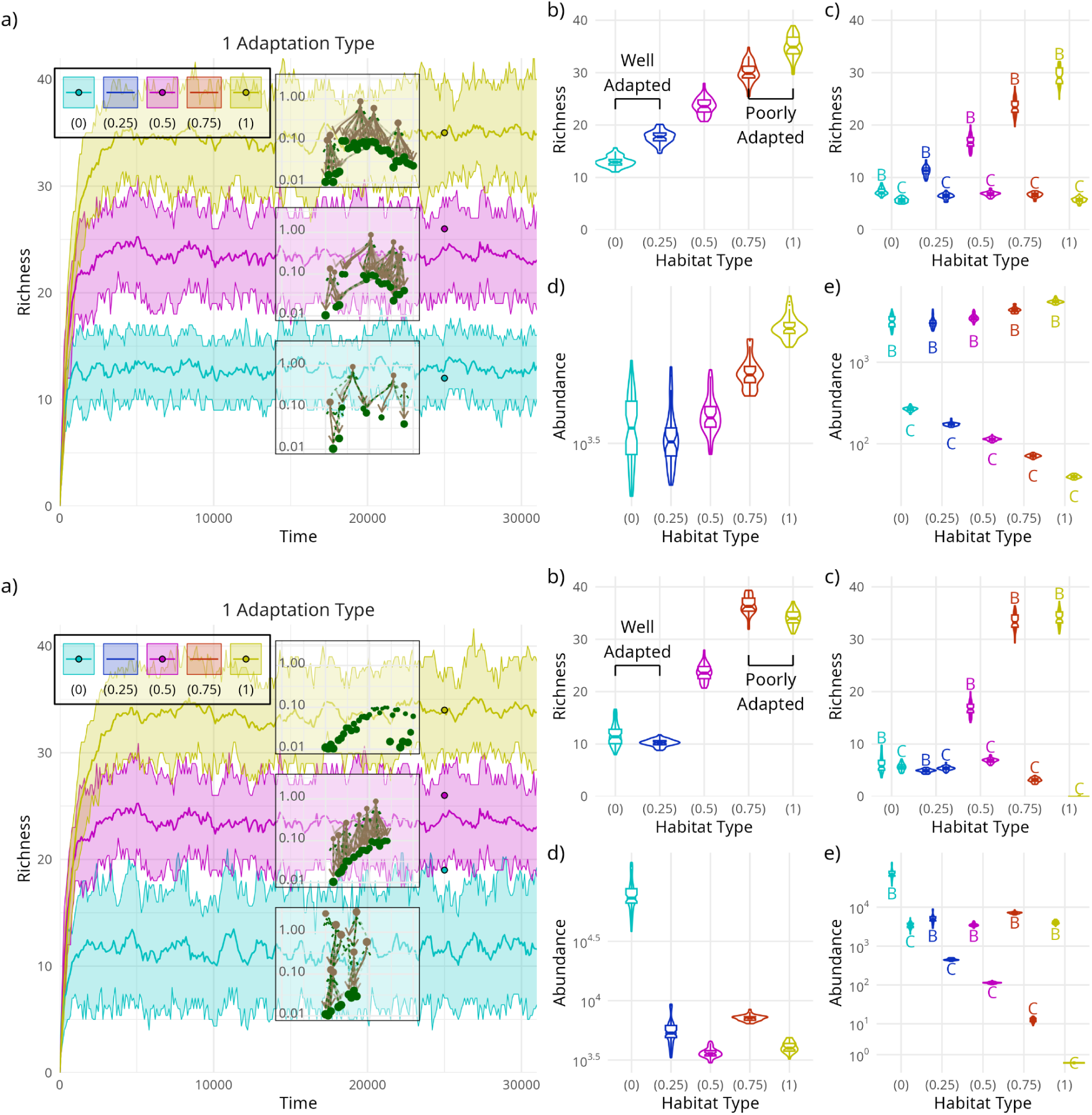
Emerging trade-offs between diversity and habitat type, contrasted when habitat-species interactions are weaker (above) or stronger (below). Panels are as described in Figure 2 of the main text. Weaker interactions use a b = 2-fold maximum effect, while stronger interactions use a b = 10-fold maximum effect; see Materials and Methods: Model for details.

**Figure S3:**
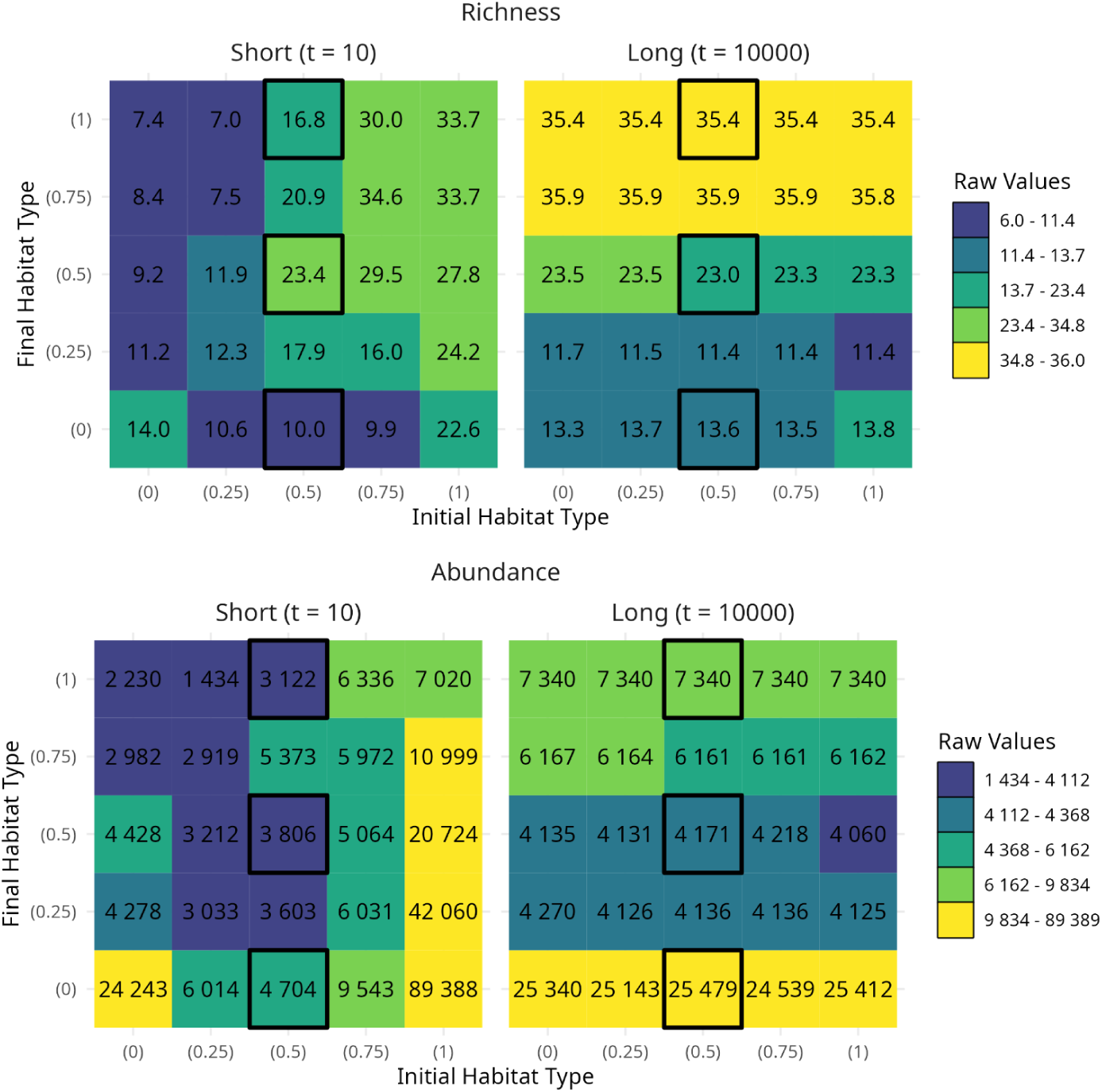
Averaged raw richness and total abundance values after (non-)intervention when there is a single adaptation type. Heatmaps show average values across the 44 systems for each combination of initial and final habitat types for richness (above) and abundance (below) across short (left, 10 time units after intervention) and long (right, 10000 time units after intervention) time scales. The short time scale analysis is for the community at 10 time units after intervention, while the long time scale analysis is for the community at 10000 time units after intervention. Black boxes indicate scenarios shown in Figure 3.

**Figure S4:**
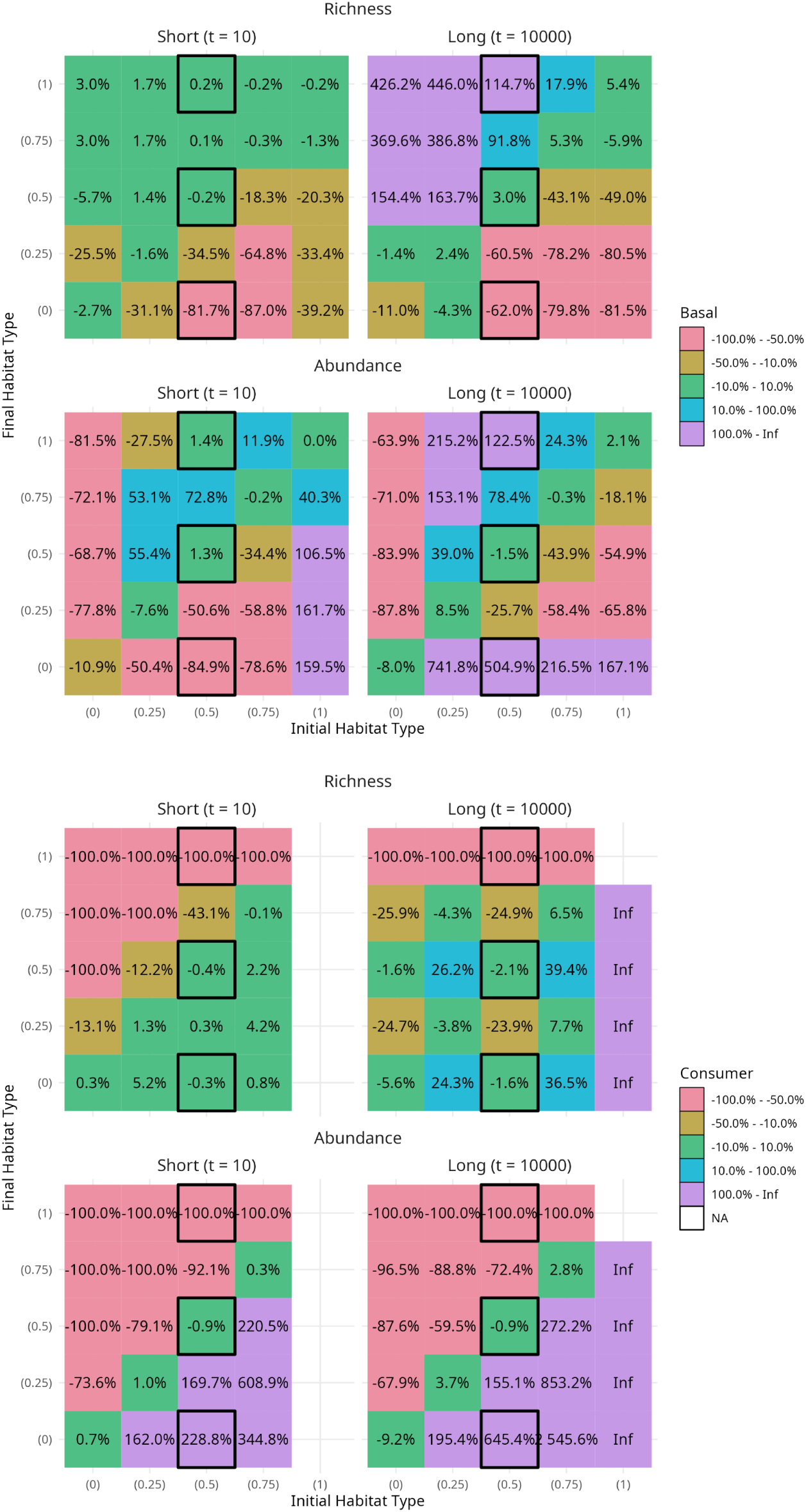
Basal (first two rows) and consumer (last two rows) contrasts between initial and final habitat types vary through time when there is a single adaptation type. Heatmaps show average percentage change (see Methods) across the 44 systems for each combination of initial and final habitat types for richness (first and third rows) and abundance (second and fourth rows) across short (left, 10 time units after intervention) and long (right, 10000 time units after intervention) time scales. The short time scale analysis compares the community at 10 time units after intervention to the community the moment just before intervention, while the long time scale analysis compares the average across 9500 - 10500 time units after intervention to the average 1000 time units before intervention. Black boxes indicate scenarios shown in Figure 3. No value is shown if there are no species before and after in any of the simulation runs (0 / 0). Infinity (Inf) is shown if there are no species before but there are species after.

**Figure S5:**
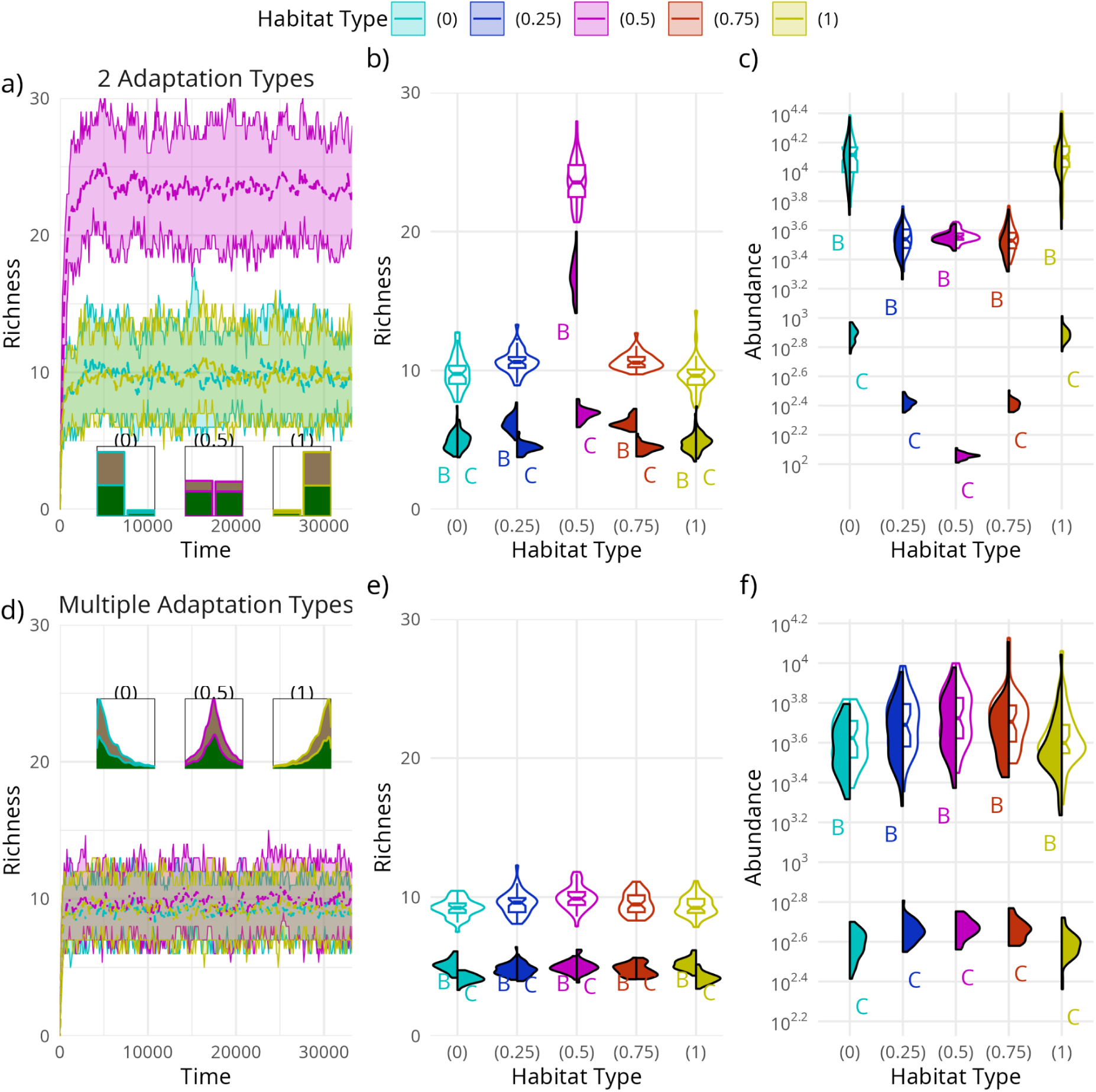
Diversity of habitat type adaptations reduces emergent richness in the local ecosystem. The first row shows results for scenarios with 2 adaptation types in the pool: half of species with adaptations to habitat type 0, half with adaptations to habitat type 1; the second row shows results for scenarios with multiple adaptation types in the pool: species adaptations range from 0 to 1. **a)** Temporal changes in species richness across 44 systems for 3 types of habitat, with lines showing the mean and ribbons the 75% interval of richness values through time (compare Figure 2a). Inset histograms capture the distribution of (either-or) adaptation traits and species type (basal, green or consumer, brown) of the emergent communities. **b)** Violin plots of richness averaged across times 20,000 and 30,000 for each simulation as a function of habitat type (compare Figure 2b). Half violins show the contributions from basal species (left, marked B) and consumer species (right, marked C). **c)** As in panel b, but for total abundance on a logarithmic scale (compare Figure 2d). Due to logarithmic scaling and the substantially larger basal abundance, consumer abundances are visually much lower. **d-f)** As in a-c), but for Multiple Adaptation Types. Insets are stacked kernel density estimates instead.

**Figure S6:**
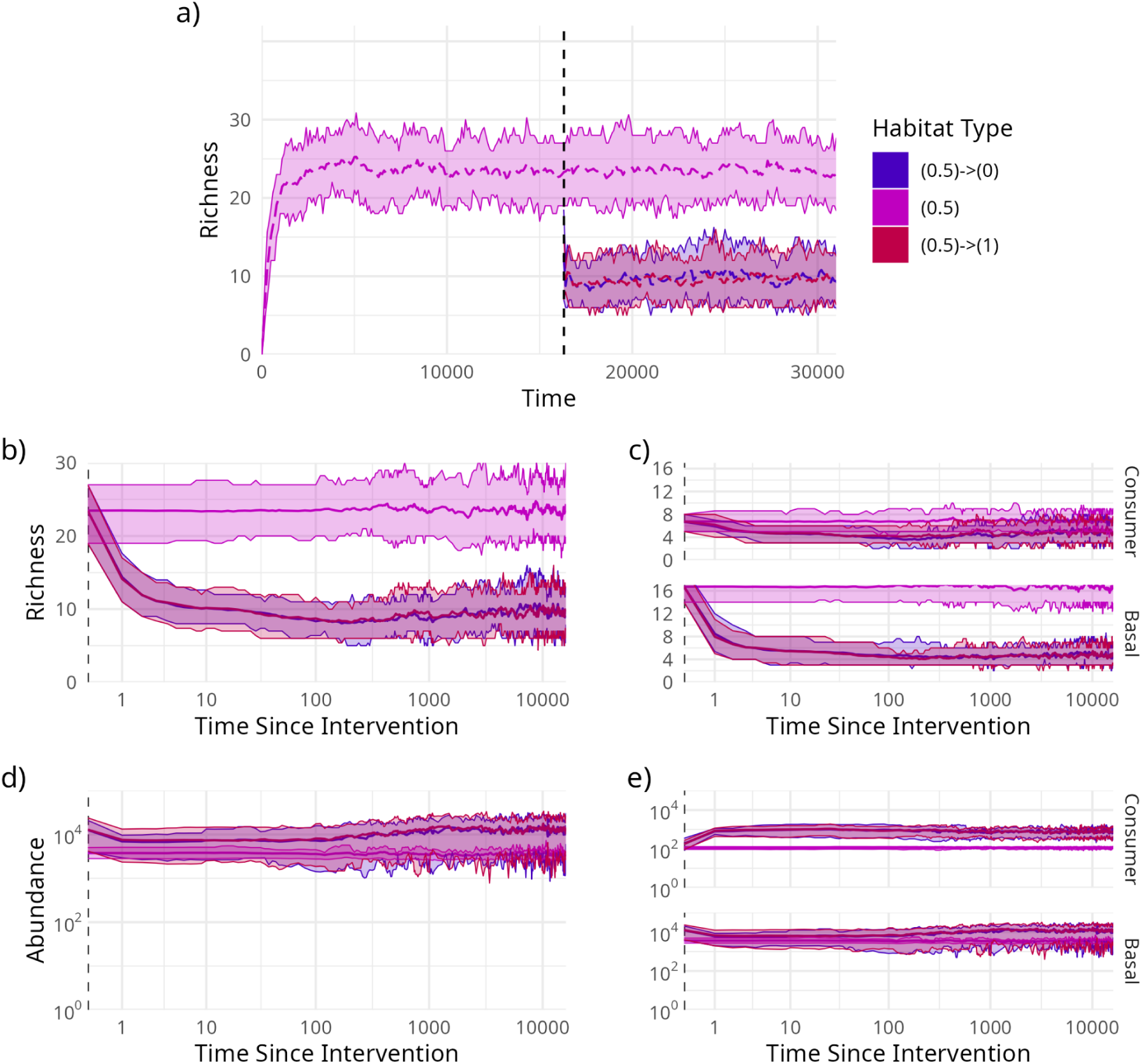
Two types of habitat type adaptations in the pool obscure changes in the structure of the system over short time scales. a) Temporal changes in species richness across 44 systems for 3 habitat change scenarios: intermediate 0.5 habitat type without intervention and with intervention to either extreme 0 or 1 habitat types. The vertical dashed black line indicates when interventions occur. b) Richness changes after the (non-)intervention on a log(1+Time) axis highlighting the difference in short term and long term post-intervention behaviour. c) As in panel b, but with richness partitioned amongst consumer (top panel) and basal (bottom panel) species. d) As in b, but for total abundance. e) As in c, but for total abundance. For all panels, solid lines show the mean across the 44 system runs, while ribbons show the range containing 75% of the richness values across the systems through time.

**Figure S7:**
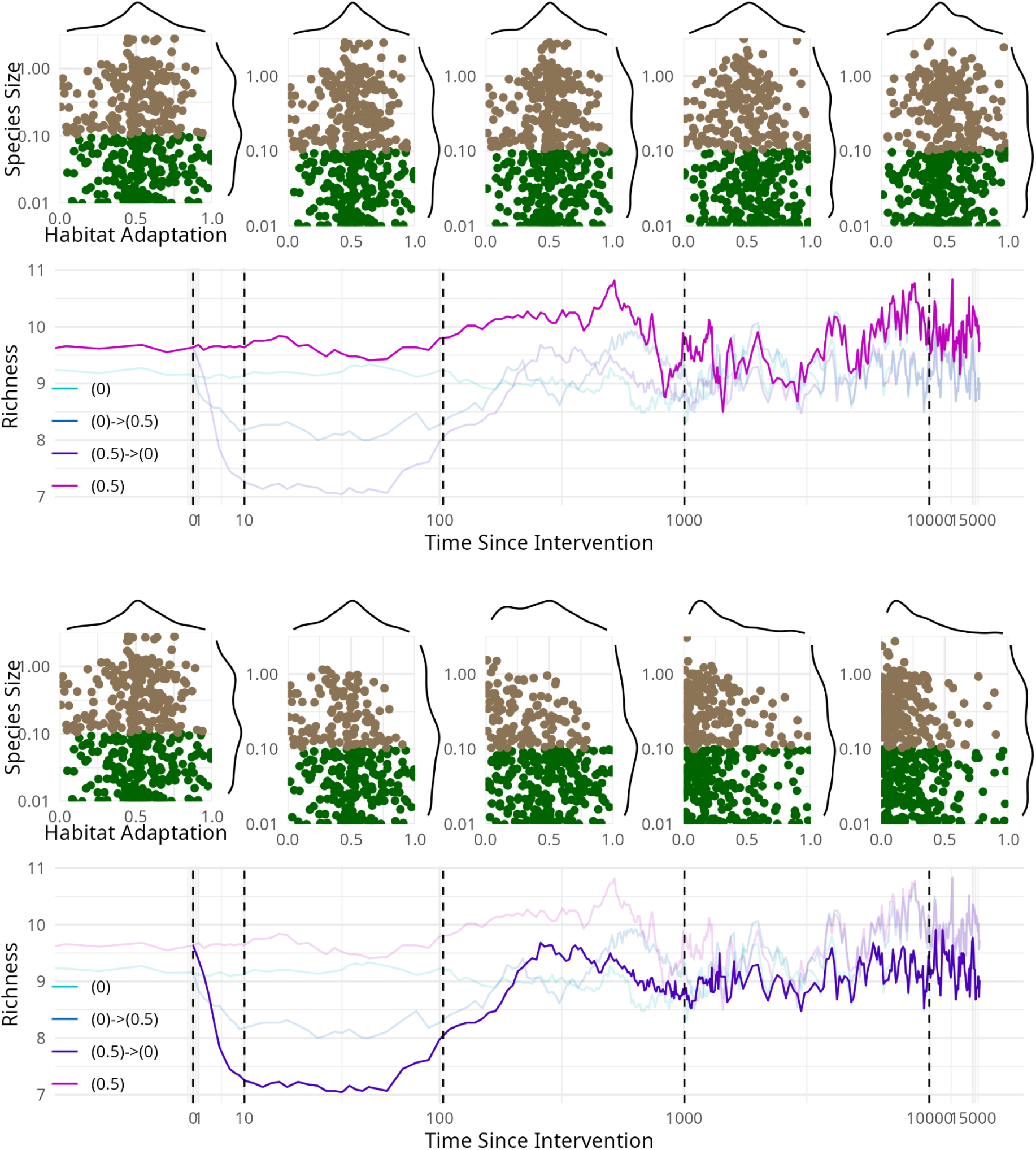
Contrasting non-intervention (first and second rows) and intervention (third and fourth rows) cases show how community structures change across time scales. First row: non-intervention habitat type 0.5 plots of points and corresponding marginal densities – of species size against habitat adaptation type – representing each species present in each of 44 communities at the times corresponding to the horizontal lines in the second row. Second row: the mean richness through time of the 44 system runs for select example (non-)interventions, with the 0.5 emphasised. Third row: as in the first row, but for the intervention from habitat type 0.5 to 0. Fourth row: as in the second row, but for the intervention from habitat type 0.5 to 0.

**Figure S8:**
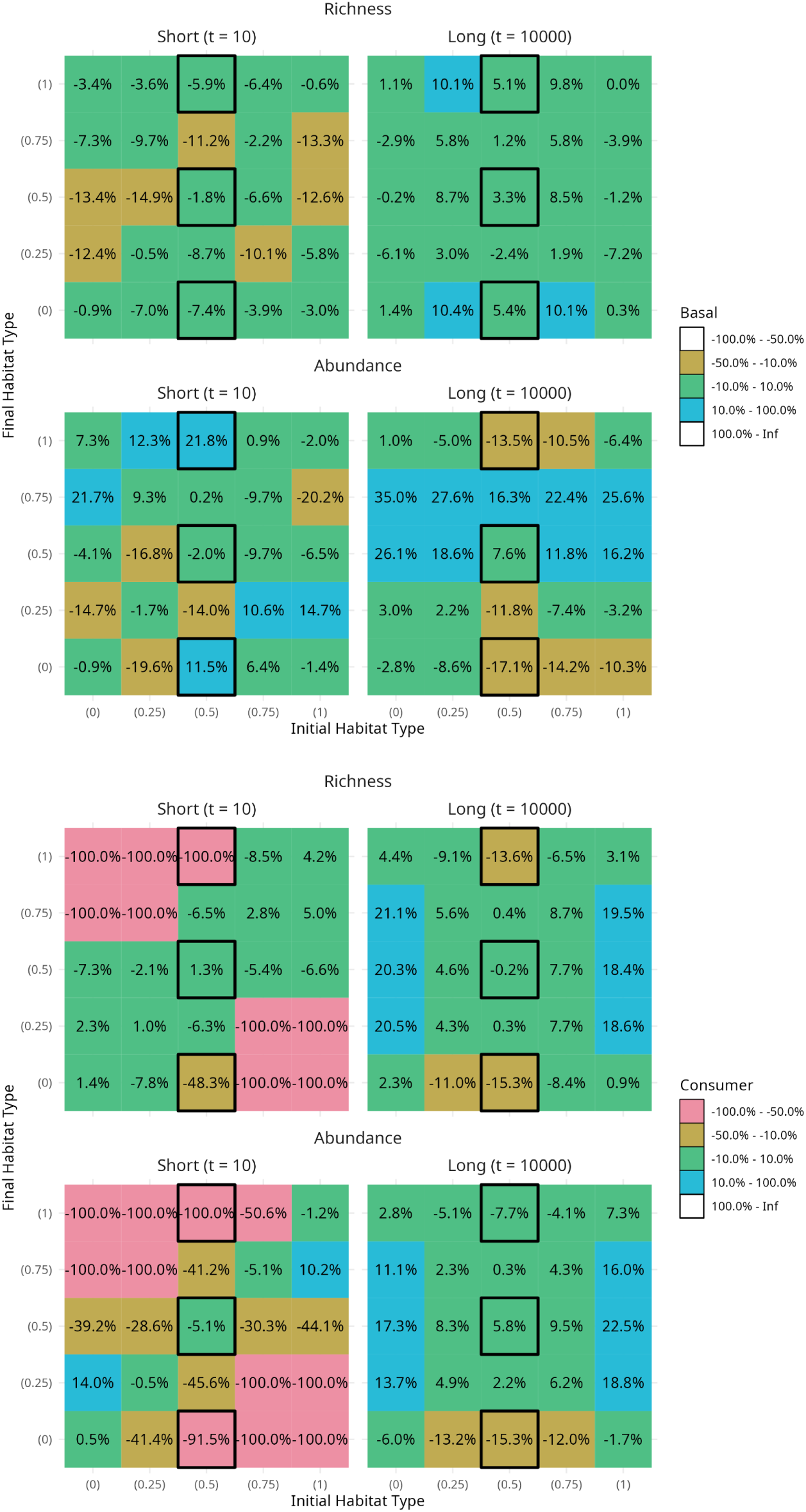
Basal (first two rows) and consumer (last two rows) contrasts between initial and final habitat types vary through time when there are multiple adaptation types. Heatmaps show average percentage change (see Methods) across the 44 systems for each combination of initial and final habitat types for richness (first and third rows) and abundance (second and fourth rows) across short (left, 10 time units after intervention) and long (right, 10000 time units after intervention) time scales. The short time scale analysis compares the community at 10 time units after intervention to the community the moment just before intervention, while the long time scale analysis compares the average across 9500 - 10500 time units after intervention to the average 1000 time units before intervention. Black boxes indicate scenarios shown in Figure 3.

**Figure S9:**
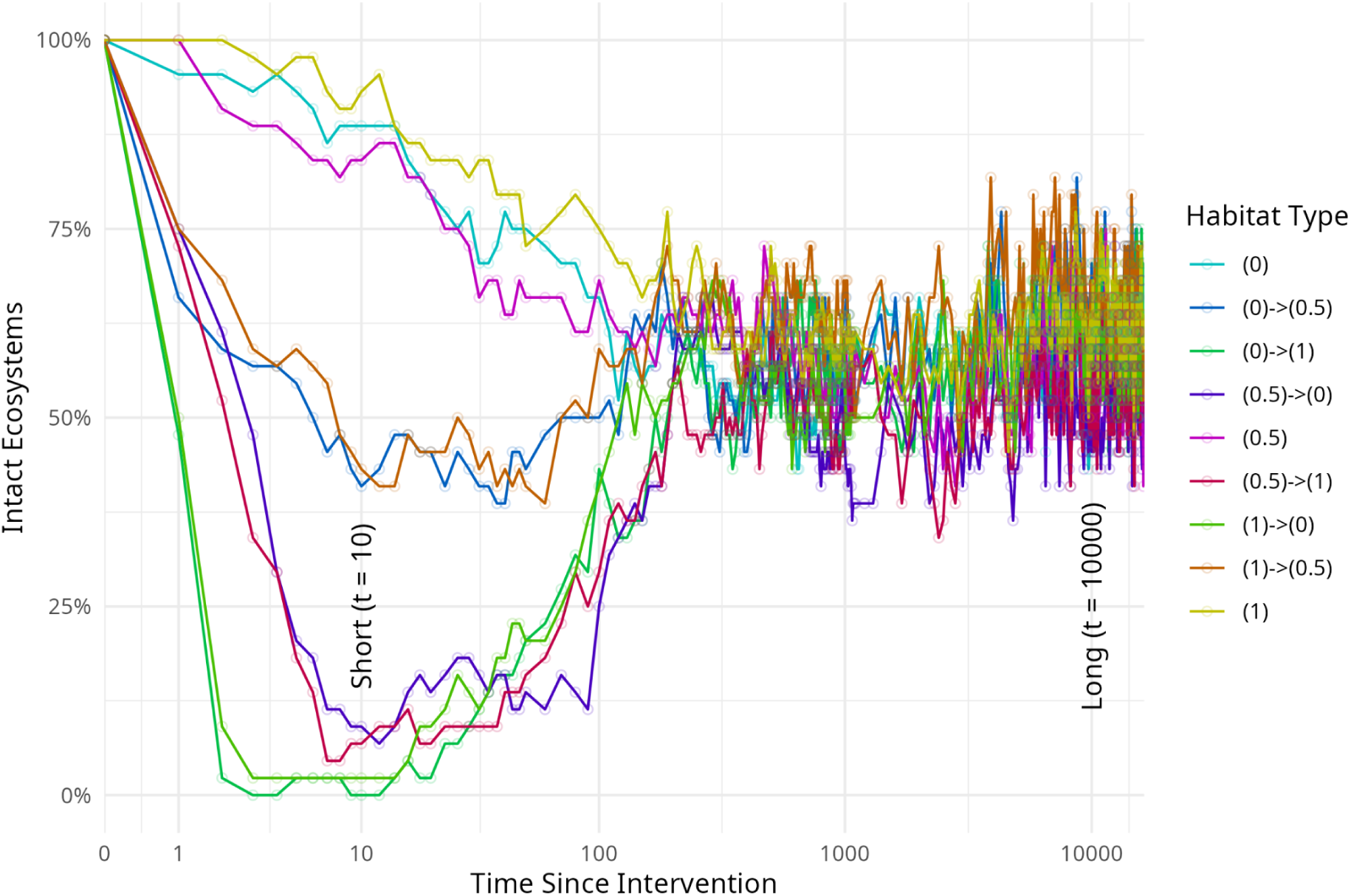
Ecosystems experiencing larger interventions to habitat type lose their intact status more quickly and more often than systems with smaller or no-intervention. We plot the percentage of the 44 systems for each intervention scenario that remain intact – here, have non-negative net richness change – through time on a Log(1+Time) axis when there are multiple adaptation types. Note that the slightly higher richness of the 0.5 final habitat type (see Figure 5) creates additional clustering of the different intervention scenario lines.

**Figure S10:**
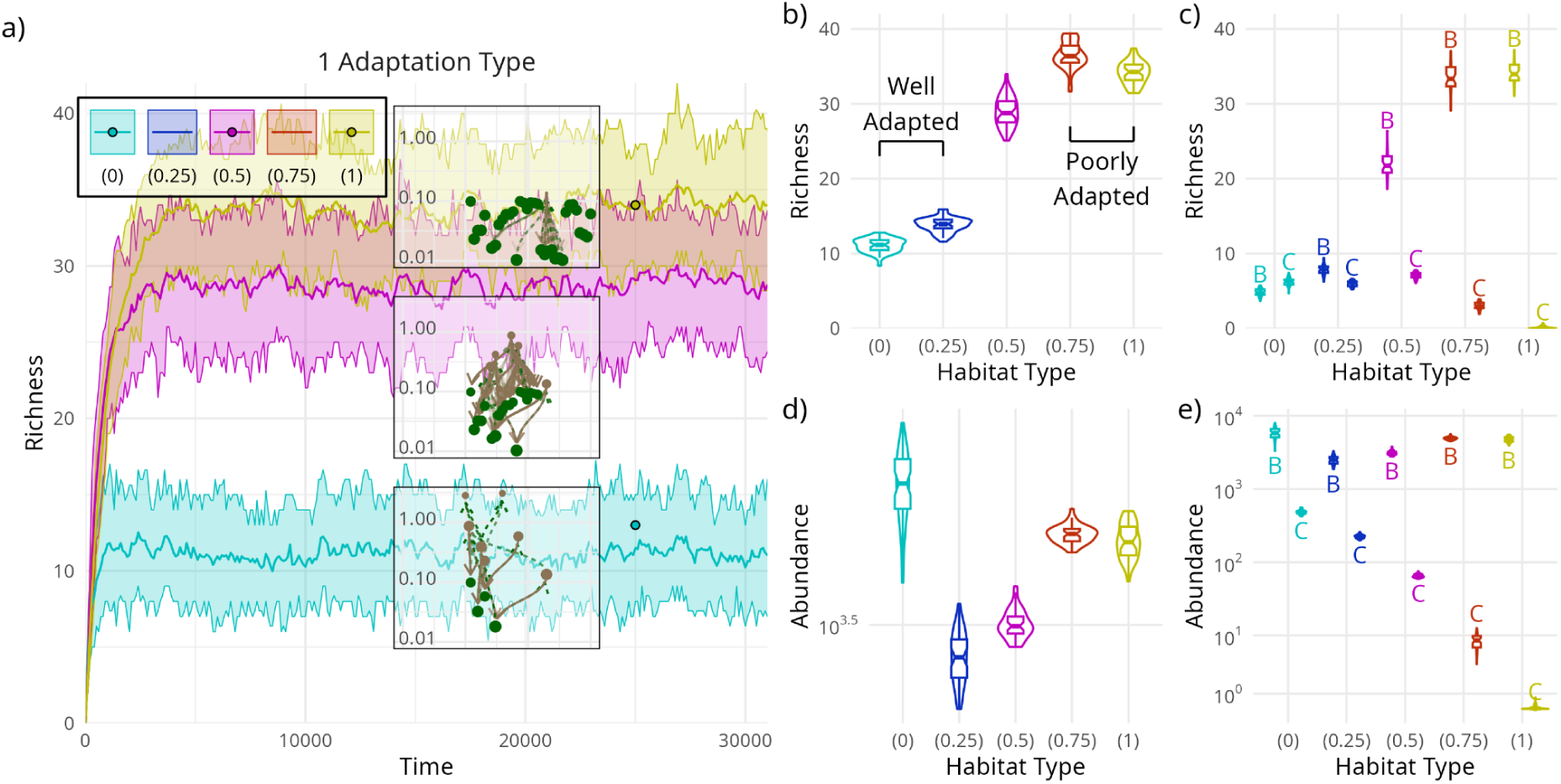
Results for scenarios with a single adaptation type in the pool are not significantly altered due to the presence of basal interspecific competition. a) Temporal changes in species richness across 44 systems for 3 types of habitat (intermediate scenarios are omitted for simplicity) in the absence of habitat interventions. Different colours represent different habitat scenarios (e.g. yellow: the habitat is of type 1). For a given scenario, solid lines show the mean across the 44 system runs, while ribbons show the range containing 75% of the richness values across the systems through time. Points represent one system evaluated at the same time for each scenario. Insets show how resultant communities, points in a, are structured for the different scenarios (brown = consumer species, green = basal species, with lines connecting consumer and consumed species). b) Violin plots of richness averaged across times 20,000 and 30,000 for each simulation as a function of habitat type. Note that species adaptation to the habitat is highest in the left hand side scenarios, and lowest in the right hand ones. c) As in panel b, but with richness displayed by whether the species are basal species (violins marked B) or consumer species (violins marked C). d-e) As in panels b and c, but for total abundance (i.e., abundance summed over all species in panel d or over all basal or consumer species in panel e).

**Figure S11:**
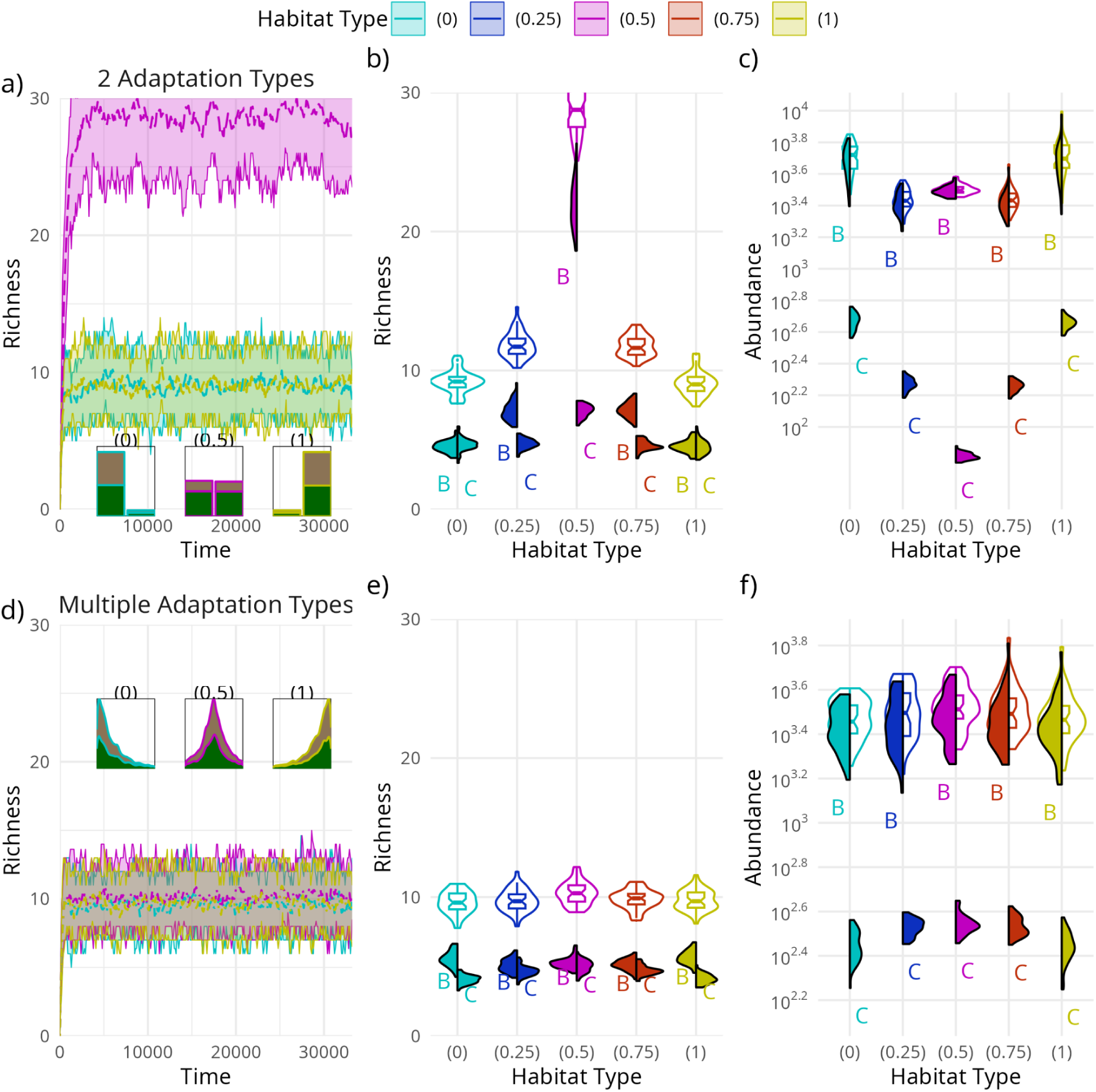
Results for a diversity of habitat type adaptations are not significantly altered due to the presence of basal interspecific competition. The first row shows results for scenarios with 2 adaptation types in the pool: half of species with adaptations to habitat type 0, half with adaptations to habitat type 1; the second row shows results for scenarios with multiple adaptation types in the pool: species adaptations range from 0 to 1. a) Temporal changes in species richness across 44 systems for 3 types of habitat, with lines showing the mean and ribbons the 75% interval of richness values through time (compare Figure 2a). Inset histograms capture the distribution of (either-or) adaptation traits and species type (basal, green or consumer, brown) of the emergent communities. b) Violin plots of richness averaged across times 20,000 and 30,000 for each simulation as a function of habitat type (compare Figure 2b). Half violins show the contributions from basal species (left, marked B) and consumer species (right, marked C). c) As in panel b, but for total abundance on a logarithmic scale (compare Figure 2d). Due to logarithmic scaling and the substantially larger basal abundance, consumer abundances are visually much lower. d-f) As in a-c), but for Multiple Adaptation Types. Insets are stacked kernel density estimates instead.

